# Segregation and integration of the functional connectome in neurodevelopmentally ‘at risk’ children

**DOI:** 10.1101/2021.05.11.443579

**Authors:** J. S. Jones, the CALM Team, D. E. Astle

## Abstract

Functional connectivity within and between Intrinsic Connectivity Networks (ICNs) transforms over development and supports high order cognitive functions. But how variable is this process, and does it diverge with altered cognitive developmental trajectories? We investigated age-related changes in integration and segregation within and between ICNs in neurodevelopmentally ‘at-risk’ children, identified by practitioners as experiencing cognitive difficulties in attention, learning, language, or memory. In our analysis we used performance on a battery of 10 cognitive tasks, alongside resting-state functional Magnetic Resonance Imaging in 175 at-risk children and 62 comparison children aged 5-16. We observed significant age-by-group interactions in functional connectivity between two network pairs. Integration between the ventral attention and visual networks and segregation of the limbic and fronto-parietal networks increased with age in our comparison sample, relative to at-risk children. Furthermore, functional connectivity between the ventral attention and visual networks in comparison children significantly mediated age-related improvements in executive function, compared to at-risk children. We conclude that integration between ICNs show divergent neurodevelopmental trends in the broad population of children experiencing cognitive difficulties, and that these differences in functional brain organisation may partly explain the pervasive cognitive difficulties within this group over childhood and adolescence.

**Research Highlights:**

- We investigated functional brain organisation and its development in 175 children who experience neurodevelopmental difficulties in cognition and behaviour, relative to a comparison sample (n=62)
- We replicated common neurodevelopmental trends across the samples: functional connectivity increased within Intrinsic Connectivity Networks and the default-mode network increasingly segregated with age
- Neurodevelopmentally at-risk children also showed different age-related changes in functional connectivity between the ventral attention and visual networks and between the fronto-parietal and limbic networks
- Furthermore, the integration between the ventral attention and visual networks in comparison children mediated age-related changes in cognition, relative to at-risk children

## Introduction

The human connectome is a complex network optimised to minimise wiring cost whilst maximising efficient communication (Bullmore and Sporns, 2012). Long distance connections between brain regions must confer significant value to warrant the substantial wiring cost. Over development this trade-off is negotiated, between the topological value of long-range connections versus the cost of forming these connections (Akarca et a. 2021). When they do form, these costly connections allow for the integration of neuronal populations over large anatomical distances, enabling complex cognitive processing and coordinated goal-directed behaviour (Johnson and Munakata, 2005). This process is mirrored by segregation. As longer connections are added to the connectome, the relative importance of local connections is diminished (Durston et al., 2006) and networks become specialised. This integration and segregation can be seen in Intrinsic Connectivity Networks (ICNs), which consist of spatially distributed regions of the brain that are highly co-activated and thus functionally connected. ICNs are emergent properties of resting brain activity (Barnes et al., 2016; Yeo et al., 2011), correspond to modules of the connectome, and substantially overlap with major functional brain systems identified during task performance (Power et al., 2011). Importantly, individual ICNs do not necessarily operate in isolation, and the integration (increased functional connectivity) and segregation (decreased functional connectivity) between ICNs is important for flexible cognition (Cohen and D’Esposito, 2016).

This functional topology emerges as the brain develops through childhood and adolescence, coinciding with gross structural changes in the brain (Carlson et al., 2013; Morgan et al., 2018). Over this period there are marked increases in regional specialisation and global integration, as functional connectivity between anatomically proximal regions weakens with age, while longer-range connections strengthen, forming ICNs (Fair et al., 2013; Farrant and Uddin, 2015; de Lacy and Calhoun, 2018; Satterthwaite et al., 2013; Solé-Padullés et al., 2016; Tomasi and Volkow, 2014). This widely reproduced finding suggests that ICNs become increasingly coherent with age. Between ICNs, a prominent finding is that activity in the default mode network typically becomes increasingly anti-correlated (‘segregated’) with activity in so-called ‘task-positive networks’, such as the fronto-parietal network (Barber et al., 2013; Bo et al., 2014; Chai et al., 2014; Gu et al., 2015; de Lacy and Calhoun, 2018; Sherman et al., 2014). Although integration between other ICNs has been less studied, findings suggest that integration between ICNs generally increases later in development, across adolescence and into early adulthood (Betzel et al., 2014; Marek et al., 2015). Integration between specific ICNs appears to follow distinct developmental trajectories and, notably, the ventral attention network may become increasingly integrated with other ICNs earlier in development (Marek et al., 2015).

These developmental changes in functional connectivity within and between ICNs have been linked to individual differences in cognition and cognitive development. Integration within specific ICNs is associated with a broad array of cognitive processes (Van Den Heuvel and Pol, 2010), including: executive function (Seeley et al., 2007), numerical cognition (Moeller et al., 2015), working memory (Hampson et al., 2006), and IQ (Abbott et al., 2016; Sherman et al., 2014). Between ICNs, greater segregation of the default-mode network from task-positive networks has been associated with attentional control (Barber et al., 2015), working memory (Hampson et al., 2010), and IQ (Sherman et al., 2014). Furthermore, greater integration between the ventral attention network and other ICNs has been associated with better inhibitory control in young people and moderates the effect of age on performance (Marek et al., 2015). This increased coordination between ICNs is especially important in high-order cognitive processing (Cohen and D’Esposito, 2016; Finc et al., 2017). These associations with cognition and, particularly age-related changes in cognitive ability, suggest that the emergence of ICNs and developing interactions between them may support typical cognitive development.

Differences in ICN development may themselves be risk factors for cognitive or behavioural difficulties in development. Indeed, reduced functional connectivity within the default-mode network (Nomi and Uddin, 2015; Sripada et al., 2014) and over-connectivity between the default-mode and task-positive networks has been associated with cognitive difficulties in childhood (Cai et al., 2018; Francx et al., 2015; Lin et al., 2018; Sripada et al., 2014). Segregation of the default-mode and task-positive networks are developmentally delayed in children with poorer attention performance, and those with greater performance difficulties show greater delay (Cai et al., 2018; Francx et al., 2015; see also Kessler et al., 2016; Lin et al., 2018; Mills et al., 2018; Sripada et al., 2014). Similarly, children with Attention Deficit Hyperactivity Disorder (ADHD) do not show a maturational strengthening between the ventral attention network and right fronto-parietal network compared to non-ADHD controls (de Lacy and Calhoun, 2018). However, these differences do not appear to be tied to any particular neurodevelopmental condition. In fact, differences in functional connectivity between the fronto-parietal, ventral attention and default mode networks have been implicated in multiple neurodevelopmental conditions (Menon, 2011) and associated with difficulties even in those without a diagnosis (e.g. Sripada et al., 2014). Taken together, these findings suggest that atypical ICN development is significantly associated with cognitive or behavioural difficulties in childhood. One plausible explanation is that differences in the emergence and timing of ICN development may itself put children at increased neurodevelopmental risk of these difficulties. Although it should be noted early that it is difficult to establish causality – alterations in network development could reflect different cognitive trajectories, or the relationships could be bidirectional.

In the present study, we explored ICN development in a sample that reflects the large heterogeneous population of children experiencing neurodevelopmental difficulties in cognition and behaviour (Astle et al., 2019; Bathelt, Gathercole, Butterfield, et al., 2018; Bathelt, Gathercole, Johnson, et al., 2018; Bathelt, Holmes, et al., 2018; Holmes et al., 2019, 2020; Mareva and Holmes, 2019; Siugzdaite et al., 2020). They were recruited on the basis of experiencing difficulties in attention, learning, language and / or memory, as identified by practitioners across a variety of children’s professional services. Hereafter we refer to this cohort of children as being neurodevelopmentally ‘at-risk’, referring to their broad heterogeneous nature, and the elevated likelihood that they will experience educational underachievement (Gathercole et al., 2016), underemployment (Emerson and Hatton, 2008) and mental health difficulties (Emerson and Hatton, 2007). Exploring resting functional connectivity in this large mixed sample of children at neurodevelopmental risk, we wanted to answer the following questions: *Firstly, can we replicate reported patterns of age-related changes in ICNs? Secondly, do these age-related patterns distinguish children ‘at risk’, relative to children not identified as struggling with cognition? And, thirdly, are these age-related patterns of ICN change associated with cognitive development and do these ICN-cognition relationships differ in neurodevelopmentally at-risk children, relative to their counterparts?*

## Method

### Sample Characteristics

Behavioural data were collected from 957 children and adolescents from the Centre for Attention Learning and Memory (CALM; Holmes et al., 2019). Children in the ‘at-risk’ sample were referred by educational and health practitioners for having one or more difficulties in attention, memory, language, literacy, and numeracy. The comparison sample was recruited from the same schools but were not identified as struggling in these areas. Children were excluded from the study if they had an uncorrected hearing or visual impairment, pre-existing neurological condition, a known genetic cause for their difficulties, or if they were a non-native English speaker.

Resting-state fMRI data were available for 348 children and adolescents who opted to take part in the MRI study. High motion scans (*n* = 111) were excluded from the analysis (see ‘fMRI Preprocessing’ for details). The final fMRI sample consisted of 237 children and adolescents aged 5-17 years (*M* = 10.80, *SD* = 2.16; see Figure S1 for age distribution): 175 at-risk and 62 comparison children (see Table 1 for group characteristics). The demographics of the MRI sample were comparable to the full sample (see Table S1).

**Table 1.** Group characteristics in the final fMRI sample.

Group characteristics in the final fMRI sample
| | At-risk ($n = 175$) | Comparison ($n = 62$) |
| --- | --- | --- |
| Age in years: $M$ ($SD$) | 10.72 (2.20) | 11.03 (2.05) |
| Boys: $n$ | 115 (65.71%) | 28 (45.16%) |
| Girls: $n$ | 60 (34.29%) | 34 (54.84%) |
| No diagnosis: $n$ | 109 (62.29%) | 59 (95.16%) |
| ADHD: $n$ | 35 (20%) | 1 (1.61%) |
| Suspected ADHD: $n$ | 10 (5.71%) | 0 (0%) |
| Autism: $n$ | 13 (7.43%) | 0 (0%) |
| Dyslexia: $n$ | 17 (9.71%) | 2 (3.23%) |

*Note.* Age at MRI assessment.

### Measures

Children completed a battery of 10 computerised and paper-based cognitive assessments that evaluated phonological processing, working memory, episodic memory, nonverbal reasoning, attention, and processing speed. The battery included: the Alliteration subtest of the Phonological Assessment Battery (Frederickson et al., 1997); the Children’s Test of Nonword Repetition (Gathercole et al., 1994); the Hector Cancellation/Balloon Hunt subtest of the Test of Everyday Attention for Children II (Manly et al., 2016); the Digit Recall, Dot Matrix, Backwards Digit Recall, and Mr X subtests of the Automated Working Memory Assessment (Alloway, 2007); the Following Instructions task (Gathercole et al., 2008); delayed recall of the Stories subtest on the Children’s Memory Scale (Cohen, 1997); and the Matrix Reasoning subtest of the Wechsler Abbreviated Scale of Intelligence II (Wechsler, 2011). Measures of learning were also collected for the Word Reading and Numerical Operations subtests of the Wechsler Individual Achievement Test II (Wechsler, 2005). The full protocol and details of the measures are described in Holmes et al. (2019). Summary statistics of the age-standardised cognitive and learning measures are provided in Table 2.

**Table 2.** Cognitive and learning characteristics in the final fMRI sample.

Cognitive and learning characteristics in the final fMRI sample
|  | At-risk |  | Comparison |  | ANOVA |  |  |
| --- | --- | --- | --- | --- | --- | --- | --- |
| | *n* | *M (SD)* | *n* | *M (SD)* | *F* | *p* | $\eta^2_p$ |
| AWMA Digit Recall | 174 | 94.08 (16.43) | 62 | 110.21 (15.88) | 48.50 | <0.001 | 0.17 |
| AWMA Backwards Digit Recall | 175 | 92.94 (12.25) | 62 | 108.67 (15.71) | 78.90 | <0.001 | 0.25 |
| AWMA Dot Matrix | 175 | 93.13 (14.94) | 62 | 105.18 (15.65) | 34.92 | <0.001 | 0.13 |
| AWMA Mr X | 175 | 96.08 (14.64) | 62 | 110.15 (18.56) | 42.02 | <0.001 | 0.15 |
| CMS Stories Delayed Recall | 174 | 8.24 (3.19) | 61 | 11.30 (3.36) | 39.13 | <0.001 | 0.14 |
| CNRep | 85 | 86.95 (19.90) | 62 | 98.56 (16.74) | 13.75 | <0.001 | 0.09 |
| Following Instructions | 165 | 98.71 (13.28) | 58 | 109.71 (13.35) | 30.84 | <0.001 | 0.12 |
| PhAB Alliteration | 175 | 92.94 (9.35) | 62 | 98.08 (8.21) | 17.92 | <0.001 | 0.07 |
| TEA-Ch-II Cancellation | 170 | 10.47 (3.39) | 57 | 12.40 (2.66) | 17.15 | <0.001 | 0.07 |
| WASI-II Matrix Reasoning | 175 | 44.16 (10.36) | 62 | 53.92 (8.95) | 46.10 | <0.001 | 0.16 |
| WIAT-II Word Reading | 173 | 87.63 (17.71) | 61 | 108.38 (11.69) | 76.65 | <0.001 | 0.25 |
| WIAT-II Numerical Operations | 151 | 88.89 (18.23) | 62 | 116.34 (20.32) | 107.43 | <0.001 | 0.34 |

*Note.* Age-normalised scores are reported for all tests with normative data, except for the following instructions test where scores have been age-residualised, mean-centered and scaled ( $M=100$ , $SD=15$ ). Analysis of variance tests examined group differences in age-standardised scores including gender as an effect of no interest. Automated Working Memory Assessment (AWMA), Children's Memory Scale (CMS), Children's test of Nonword Repetition (CNRep), Phonological Assessment Battery (PhAB), Test of Everyday Attention for Children II (TEA-Ch-II), Wechsler Abbreviated Scale of Intelligence II (WASI-II), Wechsler Individual Achievement Test II (WIAT-II).

The dimensionality of the raw data was reduced using Principal Components Analysis (PCA) with varimax rotation using the principal function from the psych package (version 2.0.9) in R (version 4.0.3). The raw cognitive scores from the full sample of children with behavioural data (*N* = 957) were first scaled to unit variance and mean-centred. Missing data were then imputed using K-nearest neighbours with the knn function from the impute package (K = 10, version 1.64.0). Across all variables and participants, 6.02% of data were missing and imputed (see Tables S2-S4 for a summary). From the PCA, we extracted component scores for the first two rotated components, which explained 64.8% variance in the data. Two components were extracted because additional components were primarily defined by a high loading on only one variable and adding a third component only explained an additional 6.1% of variance in the data. The first component predominantly loaded on executive measures of working memory, non-verbal reasoning, and selective attention; whereas the second component predominantly loaded on verbal measures of phonological processing and memory (see Figure 1). The executive vs. phonological interpretation of these component loadings is consistent with a recent factor analysis in the same sample (Holmes et al., 2020). At-risk children had significantly lower scores on the executive component (*M* = 0.09, *SD* = 0.93) relative to comparison children (*M* = 1.03, *SD* = 1.17) with age and gender included as covariates, *F*(1, 233) = 52.27, *p* = 6.87 x 10^−12^, *η*^2^_p_ = 0.18. Similarly, at-risk children had significantly lower scores on the phonological component (*M* = 0.18, *SD* = 0.83) relative to comparison children (*M* = 0.73, *SD* = 0.69), *F*(1, 233) = 20.69, *p* = 8.67 x 10^−6^, *η*^2^_p_ = 0.08. Component scores were used in subsequent analyses to examine whether age-related changes in cognition are mediated by functional connectivity.

**Figure 1.**
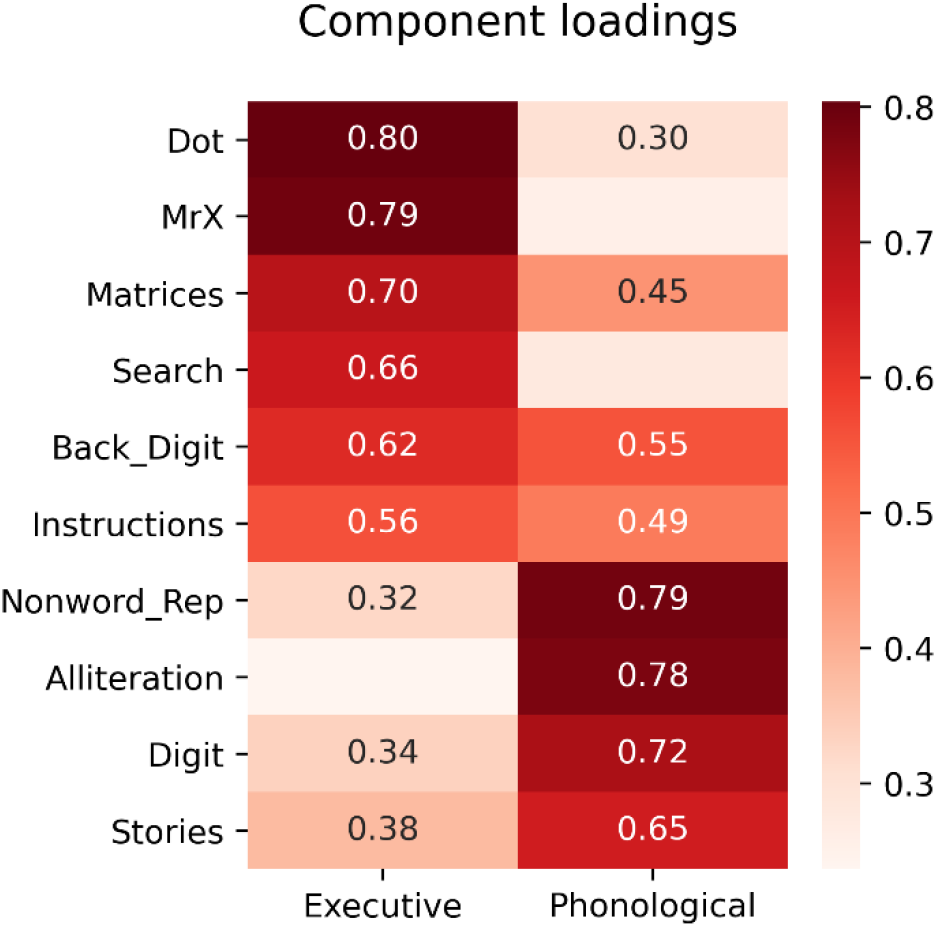
Loadings of cognitive variables on the two rotated principal components. *Note*. Loadings lower than 0.3 are suppressed for visualisation purposes. Dot (Dot Matrix), Matrices (Matrix Reasoning), Search (Hector Cancellation/Balloon Hunt), Back_Digit (Backwards Digit Recall), Instructions (Following Instructions), Nonword_Rep (Children’s test of Nonword Repetition), Digit (Digit Recall).

### Image Acquisition

Magnetic resonance imaging data were acquired at the MRC Cognition and Brain Sciences Unit, University of Cambridge. All scans were obtained on a Siemens 3T Prisma-Fit system (Siemens Healthcare, Erlangen, Germany) using a 32-channel head coil.

In the resting-state fMRI, 270 T2*-weighted whole-brain echo planar images (EPIs) were acquired over nine minutes (time repetition [TR] = 2s; time echo [TE] = 30ms; flip angle = 78 degrees, 3×3×3mm). The first 4 volumes were discarded to ensure steady state magnetization. Participants were instructed to lie still with their eyes closed and to not fall asleep. For registration of functional images, T1-weighted volume scans were acquired using a whole-brain coverage 3D Magnetization Prepared Rapid Acquisition Gradient Echo (MP RAGE) sequence acquired using 1-mm isometric image resolution (TR = 2.25s, TE = 2.99ms, flip angle = 9 degrees, 1×1×1mm).

### fMRI Pre-processing

Available resting-state fMRI data from 348 children was minimally pre-processed in fMRIPrep version 1.5.0 (Esteban et al., 2019), which implements slice-timing correction, rigid-body realignment, boundary-based co-registration to the structural T1, segmentation, and normalisation to the MNI template. The data were then smoothed by 6mm full-width at half-maximum. Many methods exist to denoise motion and physiological artefacts from resting-state fMRI; however, the effectiveness of these strategies varies depending on the sample (Ciric et al., 2017; Parkes et al., 2018). We evaluated the performance of several denoising strategies (head movement regressors, aCompCor, ICA-AROMA, motion spike regression, white matter [WM] and cerebrospinal fluid [CSF] regression, and global signal regression) on several quality control metrics (edge weight density, motion-functional connectivity correlation, distance-dependence, and functional degrees of freedom lost) using the fmridenoise package in Python (Finc et al., 2019; see Supplementary materials). The most effective confound regression procedure included a band-pass filter between 0.01-0.1Hz, 10 aCompCor components from the WM and CSF signal (Behzadi et al., 2007), linear and quadratic trends, and motion spikes (framewise displacement >0.5mm; Power et al., 2012). Simultaneous confound regression was performed in the Nipype (version 1.2.0) implementation of AFNI’s 3dTproject (Cox, 1996). Children were first excluded for high average motion (mean framewise displacement >0.5mm, *n*=93) and then for a large number of motion spikes (>20% spikes, *n*=18), where few temporal degrees of freedom would have remained. The final functional connectome sample included 237 children (at-risk *n*=175, comparison *n*=62) with mean framewise displacement 0.20mm (*SD*=0.09mm).

### Network Functional Connectivity

The denoised fMRI data were parcellated into 100 cortical regions that were assigned to seven ICNs (Schaefer et al., 2018). Pearson correlations were computed for the regional time-series within each individual generating 100×100 connectivity matrices. Edge weights were transformed using Fisher’s z-transformation. Positive and negative connectomes were generated for each individual by thresholding the connectivity matrices to retain the top 25% of positive or negative edges at the group level (see Figure S7). This was done to exclude false positive edges and to ensure that the same edges across individuals are retained for comparison in subsequent analyses, as in Baum et al. (2017). To test the robustness of brain-behaviour results, connectomes were generated at additional cost thresholds (1% intervals between 15-35%). Average functional connectivity was calculated within and between seven pre-defined ICNs: visual, somatomotor, dorsal attention, ventral attention, fronto-parietal, default mode, and limbic (Yeo et al., 2011). Finally, global intra- and inter-network functional connectivity were calculated by averaging these values within and between networks respectively.

### Analyses

First, we examined whether age correlations with global intra- and inter-network functional connectivity aligned with previously reported trajectories in childhood development. We then tested whether group (at-risk vs. comparison) moderated these associations in linear models. Second, we examined whether age associations with functional connectivity between or within specific ICNs differed between the two groups. Multiple comparisons across network pairs were corrected for using the False Discovery Rate (FDR) Benjamini-Hochberg procedure (Benjamini and Hochberg, 1995). Third, we tested whether any of these age-related changes were associated with cognitive development. Specifically, whether functional connectivity mediated age-related changes in the cognitive components identified from the PCA, and whether this was moderated by group (see Figure 3a). Statistical significance was ascertained by computing 95% confidence intervals (CI) of the moderated mediation beta from 1000 bootstrapped estimates and by comparing this to the null hypothesis. In addition, we calculated the area under the curve (AUC) for beta estimates across connectome thresholds (15-35%) and compared this to estimates expected by chance in 10,000 samples with randomly shuffled group labels. All models including age*group interaction terms included gender, motion, and mean functional connectivity (pre-thresholding) as nuisance covariates. Linear regression (‘ols’) and mediation analyses (‘Mediation’) were conducted with statsmodels 0.12.1 in Python 3.8.6.

**Figure 2.**
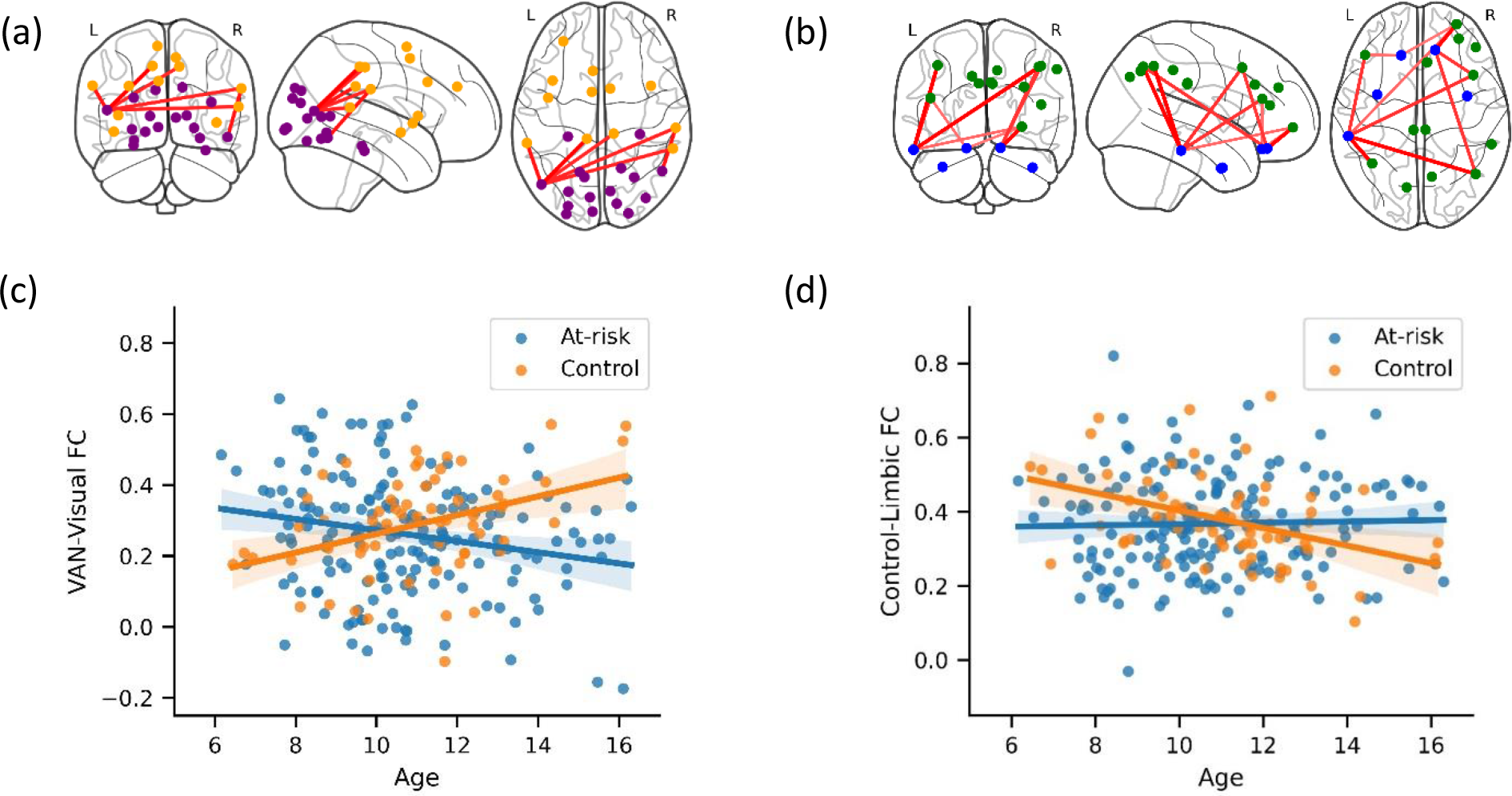
Age-by-Group interactions on functional connectivity between ICNs. *Note*. Positive edges between the visual and ventral attention networks (a) and the limbic and fronto-parietal networks (b) at 25% cost threshold. Age associations with positive functional connectivity between the visual and ventral attention networks (c) and the limbic and fronto-parietal networks (d).

**Figure 3.**
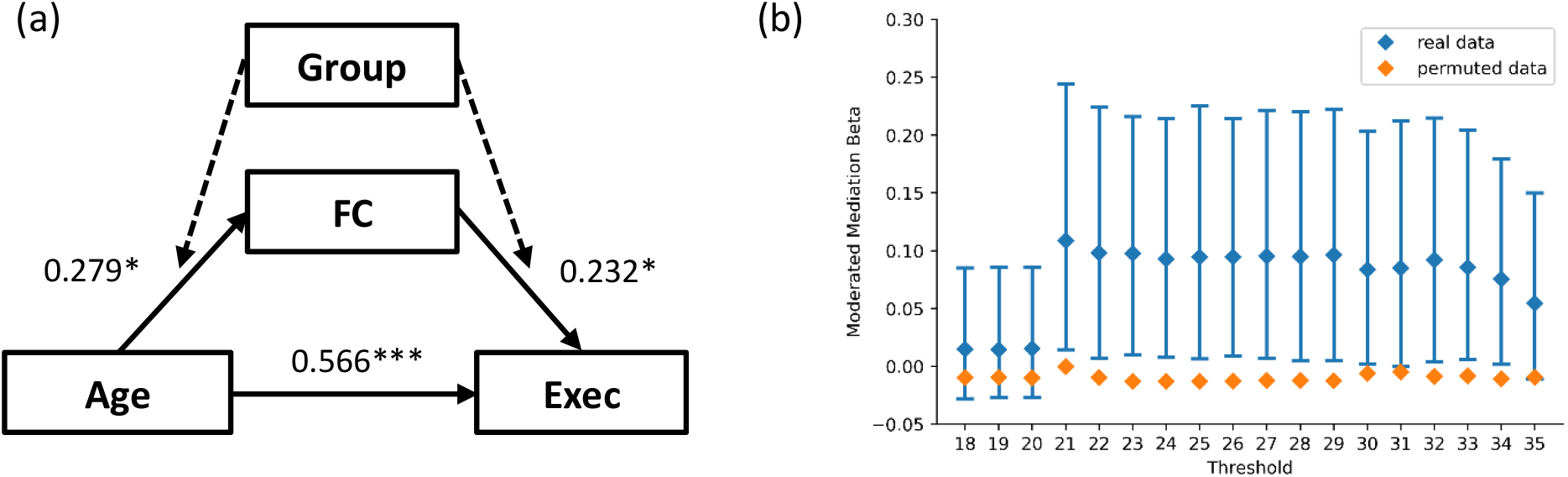
Moderated mediation of visual-ventral attention network functional connectivity on age-related changes in executive function. *Note*. (a) The moderated mediation model examining group-moderated mediation effect of positive functional connectivity (FC) between the visual and ventral attention networks on age-related changes executive function (Exec). Beta weights are shown for the control group. (b) 1000 bootstrapped estimates of the moderated mediation effect across proportional thresholds compared to permuted estimates when group labels were shuffled 10,000 times. Error bars denote 95% confidence interval. Thresholds 15-17 are not displayed because no edges were present. *p<0.05, ***p<0.001

## Results

### Age-related changes in integration and segregation

We first examined whether average functional connectivity within networks and average functional connectivity between networks correlated with age across both groups. In the positive connectome, age significantly positively correlated with average intra-network functional connectivity in the combined sample across all thresholds (*r* = 0.130-142, *p* = 0.028-045) but not with average inter-network functional connectivity (*r* = 0.016-071, *p* = 0.279-802). Age was also significantly associated with intra-network functional connectivity across all but two thresholds when controlling for gender, motion and mean functional connectivity (β = 0.101-111, *SE* = 0.051-054, *p* = 0.038-0.052). In the negative connectome, age was not associated with average intra-network functional connectivity, which was limited to default-mode connections (*r* = -0.066-096, *p* = 0.141-313), or average inter-network functional connectivity at any threshold (*r* = -0.037-104, *p* = 0.111-570).

Next, we investigated age associations with functional connectivity between or within specific ICNs. When considering both groups together, no positive connections between or within specific ICNs were significantly associated with age when controlling for gender, motion and mean functional connectivity. However, the negative connection (anti-correlation) between the dorsal attention and default-mode networks significantly strengthened with age across all thresholds (β = -0.172-0.207, *SE* = 0.058-0.060, *p* = 0.019-0.047 FDR-corrected). Together, these results in the combined sample replicate commonly reported findings in typical development: strengthening integration within ICNs and segregation of the default-mode and task-positive networks.

### Age-by-group interactions

Next, we investigated whether age associations with functional connectivity differed between the groups whilst controlling for gender, motion, and mean functional connectivity. Age associations with average intra- (β = -0.009, *SE* = 0.067, *p* = 0.897 FDR-corrected) and inter-network functional connectivity (β = 0.028, *SE* = 0.068, *p* = 0.686 FDR-corrected) did not significantly differ between the groups. However, significant age*group interactions were found for positive connections between the visual and ventral attention networks (β = 0.219, *SE* = 0.060, *p* = 0.005 FDR-corrected) and between the limbic and fronto-parietal networks (β = -0.198, *SE* = 0.065, *p* = 0.033 FDR-corrected). Older children in the comparison sample showed greater functional connectivity between the visual and ventral attention networks and reduced connectivity between the limbic and fronto-parietal networks relative to younger children, whereas at-risk children did not show these age-related changes (see Figure 2). The interaction effects were significant across multiple cost thresholds for the visual and ventral attention networks (thresholds 21-34%, β = 0.191-219, *SE* = 0.057-061, *p* = 0.004-015 FDR-corrected) and the limbic and fronto-parietal networks (thresholds 22-34%, β = -0.190-214, *SE* = 0.064-065, *p* = 0.013-048 FDR-corrected). Information about the edges included at each threshold are presented in Tables S5 and S6. We also tested whether the area under the curve for these effects across all thresholds significantly exceeded that expected by chance when group labels were randomly shuffled 10,000 times. This too indicated a significant age*group interaction on functional connectivity between the visual and ventral attention networks (AUC = 3.32, mean permuted AUC = 0.009, *p* = 0.0002) and between the limbic and fronto-parietal networks (AUC = -3.86, mean permuted AUC = -0.015, *p* = 0.0012). Age associations with negative connections within and between specific ICNs did not significantly differ between the groups.

### Links with cognition

Finally, we examined whether these age-by-group interactions predicted cognitive ability. Specifically, we examined whether age-related changes in comparison children’s executive function were mediated by functional connectivity between the visual and ventral attention networks, relative to at-risk children. Indeed, group significantly moderated the mediation effect of functional connectivity on age-related changes in executive function (β = 0.095, 95% CI [0.008, 0.214], *p* = 0.03; see Figure 3), such that the partial mediation effect was larger in comparison children, relative to at-risk children (difference in proportion of effect mediated = 12.17%, 95% CI [1.02, 30.94]). The moderated mediation was significant across multiple cost thresholds (21-34%; see Figure 3). Further, the area under the curve across all thresholds (AUC = 1.58) significantly exceeded that expected by chance when group labels were randomly shuffled 10,000 times (mean permuted AUC = -0.17, *p* = 0.0004). This was also significant when only considering cost thresholds with a unique number of between network edges (AUC = 0.76, mean permuted AUC = -0.07, *p* = 0.0008). Similar results were obtained when using the first principal component from four tasks that were previously identified as measures of a latent executive component (Holmes, Guy, 2020). This moderated mediation was significant at the 25% cost threshold (β = 0.066, 95% CI [0.004, 0.160], *p* = 0.036) and jointly across all thresholds (AUC = 0.869, mean permuted AUC = -0.134, *p* = 0.0031) and across all thresholds with a different number of edges (AUC = 0.760, mean permuted AUC = -0.057, *p* = 0.005). The effect was specific to executive function and did not generalise to phonological ability at any threshold examined (β = -0.033-0.006, *p* = 0.346-1.000). Similarly, the effect was specific to functional connectivity between the visual and ventral attention networks. Functional connectivity between the limbic and fronto-parietal networks did not significantly mediate age-related changes in comparison children’s executive function (β = -0.049-0, *p* = 0.302-990) or phonological ability at any threshold (β = 0.027-0.062, *p* = 0.182-604), relative to at-risk children.

## Discussion

We investigated whether age-related changes in ICN integration and segregation differed between neurodevelopmentally at-risk children, identified by practitioners as experiencing difficulties in cognition, relative to a comparison sample recruited from the same schools, and whether this was related to cognitive development. Across the samples we replicated common neurodevelopmental trends: increasing integration within ICNs (Farrant and Uddin, 2015; de Lacy and Calhoun, 2018; Satterthwaite et al., 2013; Sherman et al., 2014; Solé-Padullés et al., 2016; Tomasi and Volkow, 2014) and segregation of the default-mode network from the dorsal attention network (Barber et al., 2013; Bo et al., 2014; Chai et al., 2014; Gu et al., 2015; de Lacy and Calhoun, 2018; Sherman et al., 2014). However, children identified as struggling in the areas of cognition, learning, language, and memory also showed significantly different age-related changes relative to comparison children. Specifically, older comparison children had greater functional connectivity between the visual and ventral attention networks and reduced functional connectivity between the limbic and fronto-parietal networks than younger children. In contrast, ‘at-risk’ children did not show these developmental trends. Importantly, these age-related changes in connectivity significantly predicted cognitive development. Specifically, functional connectivity between the visual and ventral attention networks significantly mediated age-related changes in executive function in comparison children, relative to at-risk children.

Our findings suggest that the developing integration and segregation between ICNs follows a partially altered trajectory in children and adolescents with difficulties in the domains of attention, learning, language, and memory. Specifically, at-risk children showed a lack of integration between the visual and ventral attention networks and an absence of segregation between the limbic and fronto-parietal networks with age, compared to comparison children. This corroborates with evidence that the ventral attention network typically becomes increasingly integrated with other ICNs in late childhood (Marek et al., 2015). It also confers with reports of atypical trajectories of integration and segregation between ICNs in neurodevelopmental conditions, such as autism and ADHD, which have commonly implicated the ventral attention, fronto-parietal and default-mode networks (Abbott et al., 2016; Kessler et al., 2016; de Lacy and Calhoun, 2018; Mills et al., 2018; Sripada et al., 2014). Atypical connectivity between these networks has been highlighted as a key transdiagnostic biomarker of multiple neurodevelopmental and mental health conditions (Menon, 2011). However, while these similarities have primarily been observed across studies, our findings provide direct evidence for common neurodevelopmental patterns in a large mixed sample of children who commonly experience cognitive difficulties in childhood. Specifically, this implicates the fronto-parietal and ventral attention networks, as well as ICNs associated with visual and emotion processing.

The absence of increasing integration between the visual and ventral attention networks in neurodevelopmentally at-risk children may indicate differences in functional brain organisation and cognitive development. In typically developing children the integration between these two networks was found to mediate age-related changes in executive function, compared to at-risk children. This is in line with previous work that demonstrated increasing cross-network integration of the ventral attention network in typical childhood moderates improvements in cognitive control, as measured by performance on a visual inhibitory control task (Marek et al., 2015). Notably, the task used in this previous work and the tasks that loaded most strongly on the executive component in the current study also require visual attention/processing. Both of these cognitive functions are also represented within the primary functional roles of the ventral attention network in bottom-up attention (Corbetta and Shulman, 2002; Vossel et al., 2014) and cognitive control (Dosenbach et al., 2007; Wu et al., 2021). Furthermore, regions of the visual cortex are thought to causally influence activity in the ventral attention network on tasks requiring cognitive control and visual attention (Cai et al., 2017). Therefore, whilst maturing integration of the ventral attention and visual networks may support the development of cognitive control and/or visual attention in typical development, in our large mixed sample of children with cognitive difficulties these relationships were not present, and this may contribute to enduring cognitive difficulties experienced in this group. Crucially, it is difficult to establish causality. It may well be that these neural differences reflect, rather than drive, differential trajectories of cognitive development. We cannot disentangle the directionality, but future longitudinal data may provide a means of inferring directionality or bidirectional relationships across development.

This group difference in the mediating role of ICN integration on age-related changes in cognition was specific and robust. Age-related increases in integration between the visual and ventral attention networks was specifically associated with an executive/visual component of cognition. This component loaded heavily on measures of working memory, non-verbal reasoning, and attention, which have previously been identified as measures of an executive latent variable in a recent factor analysis of the same sample (Holmes et al., 2020). On the other hand, age-related increases in ICN integration were not associated with improvements in the phonological component. This may be because phonological processing is established earlier in development, whereas executive functions show a protracted development over childhood and adolescence (Carlson et al., 2013). The effect on executive function was reproducible when extracting only the first unrotated principal component from the subset of four tasks that were previously identified as measures of executive function (Holmes et al., 2020). This demonstrates that the effect is robust to the precise rotation and composition of tasks used to generate the cognitive component scores.

We also observed atypical development of connectivity between the limbic and fronto-parietal networks in at-risk children, such that they did not segregate with age. This was not associated with the development of executive or phonological cognition; however, it may be associated with development of ‘hot’ executive function or emotion regulation (Zelazo and Carlson, 2012). Elevated levels of behavioural difficulties have been reported in neurodevelopmentally at-risk children (Bathelt, Holmes, et al., 2018) and functional connectivity in the limbic system has been associated with emotion regulation (Posner et al., 2013), emotional lability (Hulvershorn et al., 2014), temperament (Karalunas et al., 2014), aggressiveness and conduct problems (Ho et al., 2015), and depressive symptoms (Posner et al., 2014). Furthermore, impulsivity has been associated with interactions between key nodes of the limbic and fronto-parietal networks (Li et al., 2013; Zhai et al., 2015), whereby the fronto-parietal network modulates activity in the limbic network (Baumgartner et al., 2011). The increasing segregation of these networks over typical development could indicate greater down-regulation of the limbic network emanating from the fronto-parietal network; which, speculatively, may be associated with the development of hot executive function.

There are several limitations to this investigation. First, the data are cross-sectional. We studied development by measuring age effects over the group rather than tracking individuals’ development over time. Despite this, we replicated several neurodevelopmental findings from longitudinal studies in children, including: increasing intra-network functional connectivity and increasing segregation of the default-mode and dorsal attention networks (e.g. Sherman et al., 2014). Second, the at-risk sample included a greater proportion of boys compared to the comparison sample. This is consistent with the prevalence of neurodevelopmental disorders in boys and girls (Russell et al., 2014). While gender differences in functional connectivity have been observed, boys and girls do not appear to show different developmental trajectories from childhood into early adulthood (Satterthwaite et al., 2015); nevertheless gender was included as a covariate in our analyses. Third, the ICNs were based on a parcellation of adult resting-state networks. Using a standard parcellation and group-thresholding ensured that the same anatomical regions and edges were compared across individuals. However, the cortical topography of ICNs has been shown to vary between individuals and with age (Cui et al., 2020). Future work may be improved by using more functionally homogenous individualised parcellations (Cui et al., 2020; Gordon et al., 2017). Fourth, neurodevelopmentally at-risk children may show heterogeneous development of integration and segregation between ICNs. With no clear categorical distinction between at-risk children this is difficult to test in the current study. However, future work with longitudinal data could investigate whether distinct neurodevelopmental sub-groups exist according to changes in ICN integration and segregation over time, and whether this can be predicted by baseline characteristics. Fifth, we only investigated linear relationships with age, yet cognitive and brain development can be non-linear (Luna et al., 2004; Marek et al., 2015). Our linear approach is less likely to overfit, but it may oversimplify complex neurodevelopmental changes.

In summary, neurodevelopmentally at-risk children with difficulties in the domains of attention, learning, language, and memory showed different age-related changes in ICN integration and segregation compared to typically developing children. Integration between the ventral attention and visual networks in typically developing children mediated age-related changes in executive function, compared to at-risk children. The effect was specific to this component of cognition and was robust to different degrees of connectome thresholding and dimension reduction choices. We propose that the absence of increasing integration between the visual and ventral attention networks may be a marker of enduring cognitive difficulties in neurodevelopmentally at-risk children.

## Acknowledgments

The authors were supported by the Medical Research Council program grant MC-A0606-5PQ41. We would like to thank all members of the CALM Team for their help with recruitment, data collection, and data management, as well as all of the children and parents for their participation in the study. The CALM Team includes lead investigators Duncan Astle, Kate Baker, Susan Gathercole, Joni Holmes, Rogier Kievit and Tom Manly. Data collection is assisted by a team of researchers and PhD students that includes Danyal Akarca, Joe Bathelt, Marc Bennett, Giacomo Bignardi, Sarah Bishop, Erica Bottacin, Lara Bridge, Diandra Brkic, Annie Bryant, Sally Butterfield, Elizabeth Byrne, Gemma Crickmore, Edwin Dalmaijer, Fánchea Daly, Tina Emery, Laura Forde, Grace Franckel, Delia Furhmann, Andrew Gadie, Sara Gharooni, Jacalyn Guy, Erin Hawkins, Agnieszka Jaroslawska, Sara Joeghan, Amy Johnson, Jonathan Jones, Silvana Mareva, Elise Ng-Cordell, Sinead O’Brien, Cliodhna O’Leary, Joseph Rennie, Ivan Simpson-Kent, Roma Siugzdaite, Tess Smith, Stephani Uh, Maria Vedechkina, Francesca Woolgar, Natalia Zdorovtsova, Mengya Zhang. The authors wish to thank the many professionals working in children’s services in the South-East and East of England for their support, and to the children and their families for giving up their time to visit the clinic. We would also like to thank the radiographers who support the excellent paediatric scanning at the MRC CBSU.

