## Supplementary materials for "Segregation and integration of the functional connectome in neurodevelopmentally ‘at risk’ children"

**Table S1**

*Group characteristics of the full sample*

| | Struggling learners (*n*=799) | Comparison sample (*n*=158) |
|---|---|---|
| Age in years: *M* (*SD*) | 9.42 (2.29) | 10 (2.33) |
| Boys: *n* | 550 (68.8%) | 89 (56.3%) |
| Girls: *n* | 249 (31.2%) | 69 (43.7%) |
| No diagnosis: *n* | 482 (60.3%) | 155 (98.1%) |
| ADHD: *n* | 194 (24.3%) | 1 (0.6%) |
| Suspected ADHD: *n* | 57 (7.1%) | 0 (0%) |
| Autism: *n* | 56 (7%) | 0 (0%) |
| Dyslexia: *n* | 47 (5.9%) | 2 (1.3%) |

*Note*. Age at behavioural assessment, which preceded the MRI assessment by 0.33 years on average (*SD* = 0.38).

**Figure S1**

*Age distributions in the samples*

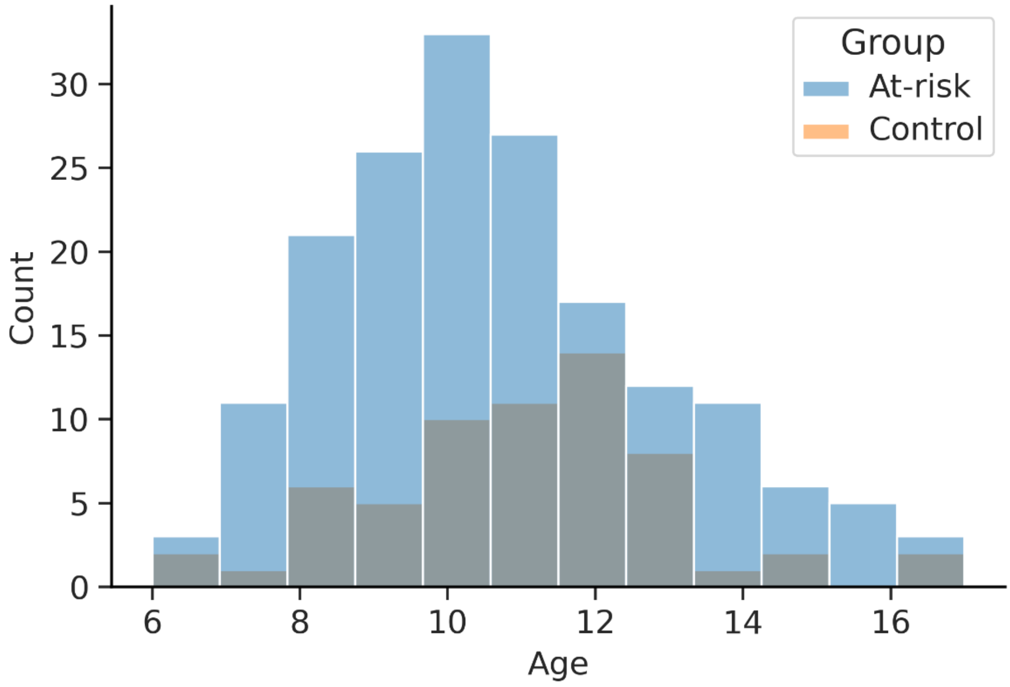


*Note*. Counts are overlaid not stacked.

**Table S2**

*Imputed data for each measure in the full sample (N=957)*

| | Missing: *n* |
|---|---|
| AWMA Backwards Digit Recall | 32 |
| AWMA Digit Recall | 6 |
| AWMA Dot Matrix | 9 |
| AWMA Mr X | 18 |
| CMS Delayed Recall | 36 |
| CNRep | 315 |
| Following Instructions | 68 |
| Matrix Reasoning | 27 |
| PhAB Alliteration | 20 |
| TEA-Ch2 Cancellation | 45 |

*Note*. The Children's test of Non-word Repetition (CNRep) was not administered to the first 300 participants. Automated Working Memory Assessment (AWMA), Children's Memory Scale (CMS), Phonological Assessment Battery (PhAB), the Test of Everyday Attention for Children 2.

**Table S3**

*Number of missing variables for each participant in the full sample (N=957)*

| Missing variables | *n* |
|---|---|
| 0 | 546 |
| 1 | 314 |
| 2 | 76 |
| 3 | 11 |
| 4 | 9 |
| 5 | 3 |
| 6 | 3 |
| 8 | 1 |

*Note*. No participants were missing 7, 9 or 10 variables

**Table S4**

*Imputed data for each measure in the MRI sample*

| | Struggling learners (*n*=175) | Controls (*n*=62) |
|---|---|---|
| AWMA Backwards Digit Recall | 0 | 0 |
| AWMA Digit Recall | 1 | 0 |
| AWMA Dot Matrix | 0 | 0 |
| AWMA Mr X | 0 | 0 |
| CMS Delayed Recall | 1 | 1 |
| CNRep | 90 | 0 |
| Following Instructions | 10 | 4 |
| Matrix Reasoning | 0 | 0 |
| PhAB Alliteration | 1 | 0 |
| TEA-Ch2 Cancellation | 5 | 5 |

### Evaluation of rsfMRI denoising strategies

A number of pipelines were evaluated to denoise motion and physiological artefacts from the resting-state fMRI data using the fmridenoise package in Python (https://github.com/compneuro-ncu/fmridenoise). The pipelines included combinations of different methods and regressors:

- 24 Head Motion Parameters (24HMP) – Regression of the 6 rigid body realignment parameters, their squares, their first derivatives, and the squares of the first derivatives
- 8 Physiological regressors (8Phys) – Regression of the time series activity from the CSF and WM masks, their squares, their first derivatives, and the squares of the first derivatives
- 4 Global Signal Regressors (4GSR) – Regression of the whole-brain time series activity, its square, first derivative, and square of the first derivative
- Motion spike regression (SpikeReg) – Regression of volumes where framewise displacement (FD) was greater than 0.5mm or where the BOLD signal change (DVARS) was 3 standard deviations away from the mean (Power, Barnes, Snyder, Schlaggar, & Petersen, 2012)
- Anatomical component correction (aCompCor) – Regression of 10 principal components estimated from the CSF and WM matter masks (Behzadi, Restom, Liau, & Liu, 2007)
- ICA-AROMA – Estimation of independent components using MELODIC in FSL, automatic identification of noise components, and their subsequent regression from the data (Pruim et al., 2015)

24HMP_aCompCor_SpikeReg (coloured purple in the following figures) was selected because it was the best performing pipeline across a number of quality control measures. It is also widely implemented as the default denoising pipeline in Conn (Whitfield-Gabrieli & Nieto-Castanon, 2012). Although GSR pipelines performed well on various quality metrics, we did not implement it here because GSR is a highly controversial technique that may remove neural signal and introduce spurious anti-correlations (Murphy & Fox, 2017).

**Figure S2**

*The density of edge weights across various denoising pipelines*

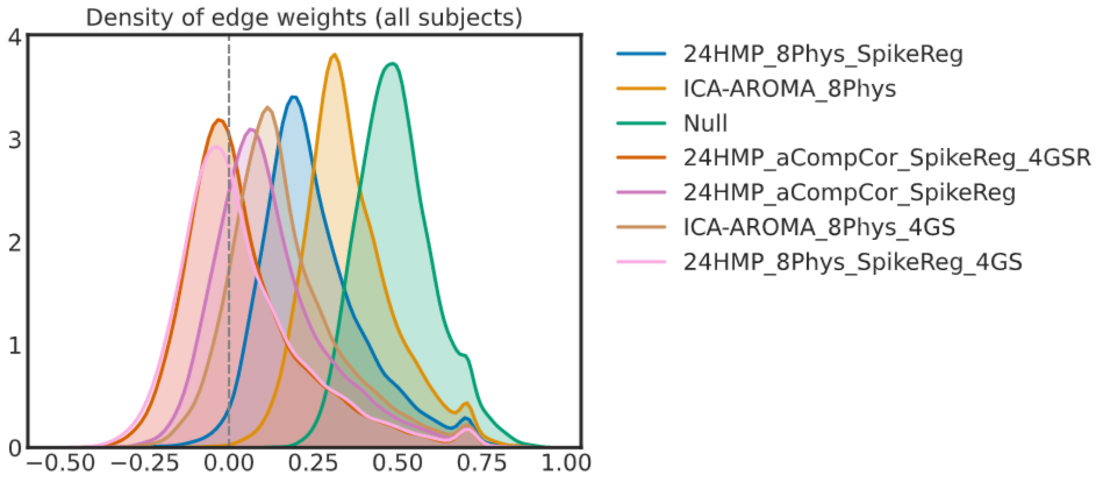

**Figure S3**

*Correlation between average movement (FD) and functional connectivity at each edge for each denoising strategy*

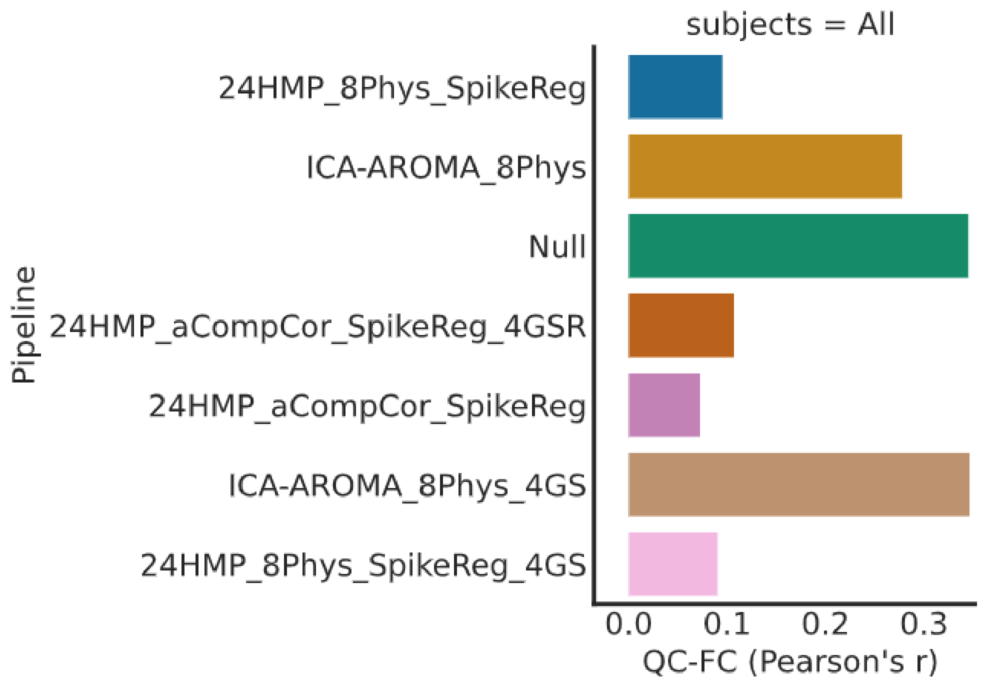


**Figure S4**

*Distance-dependence for each denoising strategy*

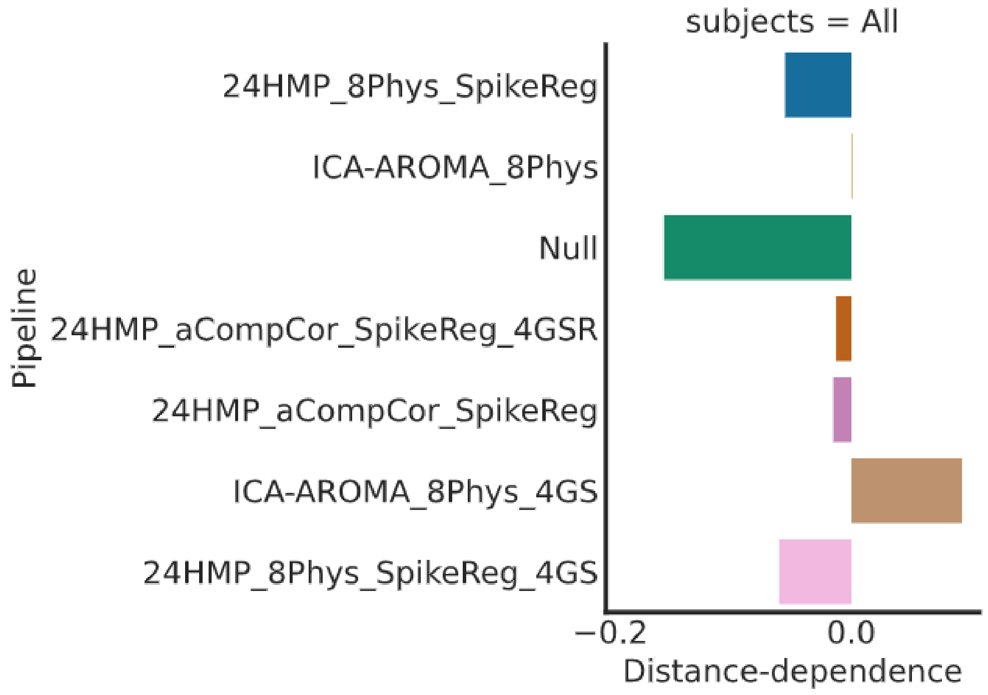


*Note.* Distance-dependence is the average correlation between functional connectivity at each edge and the Euclidean distance between nodes

**Figure S5**

*The functional degrees of freedom lost (fDOF) for each denoising strategy*

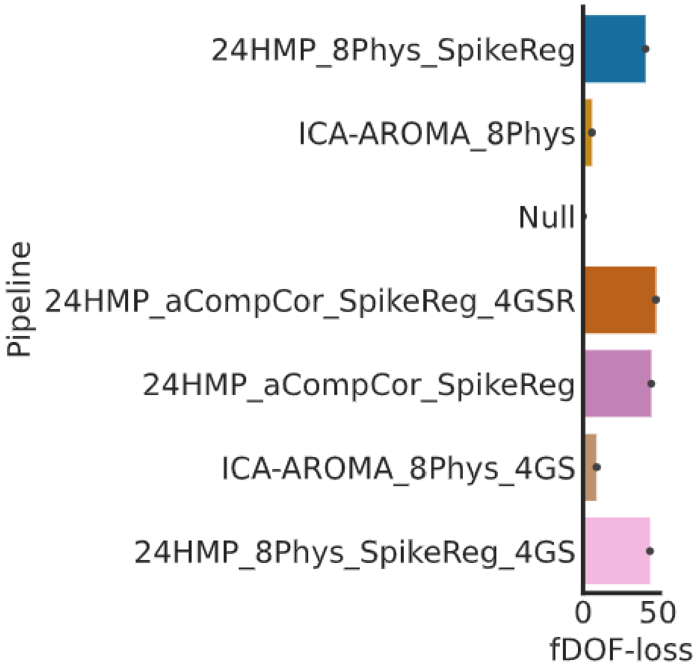


**Figure S6**

*Denoised functional connectome and correlation with motion*

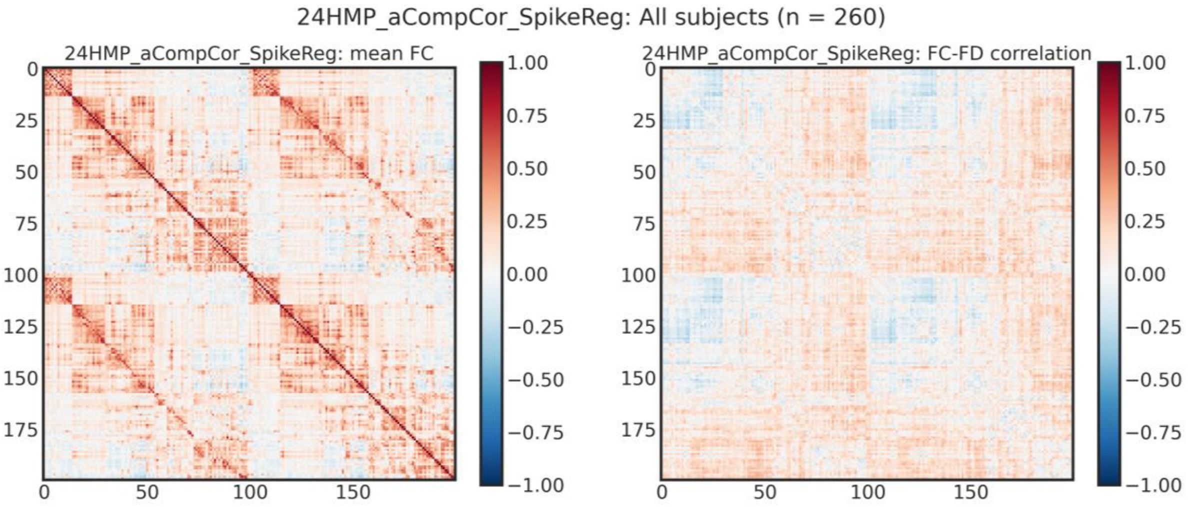


*Note*. The correlation matrix (Pearson's R) after the selected confound regression strategy including: 24 head movement parameters, 10 principal components from WM and CSF, and motion spikes (left). The corresponding correlation between average movement and functional connectivity at each edge (right). Note, the 200 region parcellation from Schaefer et al. (2018) was used here.

### Group-Thresholded Functional Connectomes

#### Figure S7

*Positive and negative group-thresholded functional connectomes.*

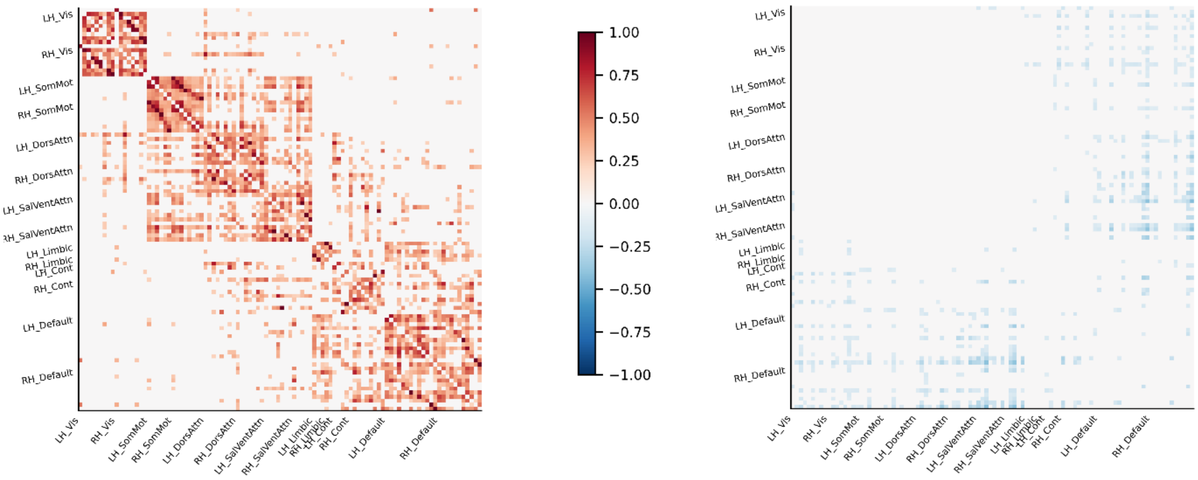


*Note.* The top 25% of positive connections (left) and the top 25% of negative connections (right) of the group-averaged functional connectome.

### Edges of the connectome between ICNs

At the 25% threshold, edges between the visual and ventral attention networks included the left visual association area MT connected to the left parietal operculum, right supramarginal gyrus, and bilateral posterior cingulate, and the right fusiform gyrus connected to the right supramarginal gyrus. Connection labels at other thresholds are displayed in Table S5.

**Table S5**

*Edges between the visual and ventral attention networks across thresholds*

| Threshold | Number of Edges | Additional edge labels at each threshold |
|---|---|---|
| 15-17 | 0 | N/a |
| 18-20 | 1 | LH_Vis_7 - LH_SalVentAttn_Med_2 |
| 21 | 3 | LH_Vis_7 - LH_SalVentAttn_ParOper_1<br>RH_Vis_3 - RH_SalVentAttn_TempOccPar_2 |
| 22 | 5 | LH_Vis_7 - RH_SalVentAttn_TempOccPar_2<br>LH_Vis_7 - RH_SalVentAttn_TempOccPar_1 |
| 23-29 | 6 | LH_Vis_7 - RH_SalVentAttn_Med_1 |
| 30 | 10 | RH_Vis_3 - RH_SalVentAttn_Med_1<br>RH_Vis_3 - LH_SalVentAttn_ParOper_1<br>RH_Vis_3 - LH_SalVentAttn_Med_3<br>RH_Vis_4 - LH_SalVentAttn_Med_3 |
| 31 | 11 | RH_Vis_3 - RH_SalVentAttn_Med_2 |
| 32 | 12 | LH_Vis_3 - RH_SalVentAttn_FrOperIns_1 |
| 33 | 14 | LH_Vis_8 - LH_SalVentAttn_Med_2<br>RH_Vis_1 - LH_SalVentAttn_FrOperIns_1 |
| 34 | 17 | LH_Vis_8 - RH_SalVentAttn_Med_1<br>LH_Vis_3 - LH_SalVentAttn_FrOperIns_2<br>RH_Vis_2 - RH_SalVentAttn_FrOperIns_1 |
| 35 | 20 | LH_Vis_2 - LH_SalVentAttn_Med_3<br>LH_Vis_5 - LH_SalVentAttn_Med_3<br>RH_Vis_3 - LH_SalVentAttn_Med_2 |

At the 25% threshold edges between the limbic and fronto-parietal network included: the left orbitofrontal cortex connected to the bilateral lateral prefrontal cortex, the right orbitofrontal cortex connected to the right lateral prefrontal cortex and right inferior parietal lobe, and the left temporal pole connected to the bilateral inferior parietal lobes and bilateral lateral prefrontal cortex. Connection labels at other thresholds are displayed in Table S6.

**Table S6**

*Edges between the limbic and fronto-parietal networks across thresholds*

| Threshold | Number of Edges | Additional edge labels at each threshold |
|---|---|---|
| 15-19 | 7 | LH_Limbic_TempPole_2 - LH_Cont_Par_1<br>LH_Limbic_TempPole_2 - LH_Cont_PFCl_1<br>LH_Limbic_TempPole_2 - RH_Cont_Par_2<br>LH_Limbic_TempPole_2 - RH_Cont_PFCl_4<br>RH_Limbic_OFC_1 - RH_Cont_Par_2<br>RH_Limbic_OFC_1 - RH_Cont_PFCl_1<br>RH_Limbic_OFC_1 - RH_Cont_PFCl_4 |
| 20 | 8 | LH_Limbic_TempPole_2 - RH_Cont_PFCl_1 |
| 21 | 9 | LH_Limbic_OFC_1 - LH_Cont_PFCl_1 |
| 22-25 | 10 | LH_Limbic_OFC_1 - RH_Cont_PFCl_1 |
| 26 | 11 | LH_Limbic_TempPole_2 - LH_Cont_Cing_1 |
| 27-29 | 13 | LH_Limbic_OFC_1 - LH_Cont_Par_1<br>LH_Limbic_OFC_1 - RH_Cont_Par_2 |
| 30 | 17 | LH_Limbic_OFC_1 - LH_Cont_Cing_1<br>LH_Limbic_TempPole_2 - RH_Cont_Cing_1<br>RH_Limbic_OFC_1 - RH_Cont_Cing_1<br>RH_Limbic_TempPole_1 - RH_Cont_PFCl_4 |
| 31 | 18 | LH_Limbic_OFC_1 - RH_Cont_PFCl_4 |
| 32-34 | 19 | RH_Limbic_OFC_1 - LH_Cont_Par_1 |
| 35 | 22 | RH_Limbic_OFC_1 - LH_Cont_PFCl_1<br>RH_Limbic_OFC_1 - LH_Cont_Cing_1<br>RH_Limbic_OFC_1 - RH_Cont_PFCl_2 |
